# Distinct neural mechanisms for the prosocial and rewarding properties of MDMA

**DOI:** 10.1101/659466

**Authors:** Boris D. Heifets, Juliana S. Salgado, Madison D. Taylor, Paul Hoerbelt, Daniel F. Cardozo Pinto, Elizabeth E. Steinberg, Jessica J. Walsh, Ji Y. Sze, Robert C. Malenka

## Abstract

The extensively abused recreational drug MDMA has shown promise as an adjunct to psychotherapy for treatment-resistant psychiatric disease. It is unknown, however, whether the mechanisms underlying its prosocial therapeutic effects and abuse potential are distinct. We demonstrate in mice that MDMA acting at the serotonin transporter within the nucleus accumbens is necessary and sufficient for MDMA’s prosocial effect. MDMA’s acute rewarding properties, in contrast, require dopaminergic signaling. MDMA’s prosocial effect requires 5-HT1b receptor activation and is mimicked by d-fenfluramine, a selective serotonin-releasing compound. By dissociating the mechanisms of MDMA’s prosocial effects from its addictive properties, we provide evidence for a conserved neuronal pathway, which can be leveraged to develop novel therapeutics with limited abuse liability.

**One Sentence Summary:** MDMA, which has both therapeutic and abuse potential, engages a brain region-specific serotonergic pathway to produce its prosocial effect.

## Main Text

MDMA is an extensively used and often abused drug that has addictive liability and toxic side effects (*1*). Yet at modest doses it has the well-documented and potentially therapeutic effects of enhancing feelings of trust, emotional openness and perhaps empathy (*2–4*). Despite the potential negative consequences of MDMA ingestion, because of its profound prosocial effects, MDMA is being evaluated to determine its efficacy in the treatment of post-traumatic stress disorder and autism spectrum disorders (*4, 5*). However, it is unknown whether the mechanisms underlying MDMA’s prosocial, therapeutic effect and abuse potential can be separated (*1*); a topic with important implications for the future therapeutic use of MDMA as well as for the development of similar agents with less potential morbidity.

MDMA is a substituted amphetamine with high affinity for the serotonin (5-HT) and dopamine (DA) transporters (SERT, DAT) (*6, 7*), through which it stimulates release of these neurotransmitters. MDMA’s acute reinforcing effects, which strongly predict addictive liability (*8, 9*), have been linked to its DA-releasing properties (*10–12*) while the role of SERT in this action is uncertain (*13, 14*). Studies on the detailed mechanisms underlying the prosocial effects of MDMA are confusing and implicate several different neuromodulatory substances including 5-HT, DA and oxytocin (Oxt) (*1, 3, 12*). Recent work highlights the role of 5-HTergic signaling in social approach behaviors, particularly within the nucleus accumbens (NAc) (*15, 16*), a conserved brain region that regulates appetitive behavior. We therefore hypothesized that MDMA’s interaction with SERT specifically in the NAc fully accounts for MDMA’s prosocial effect, but not its rewarding effect.

To study the prosocial effects of MDMA, we used the three-chamber social approach assay (*17*) (**Fig. 1A**). We found that MDMA dose-dependently increased the time a free mouse explores the chamber containing an age and sex-matched conspecific stranger mouse kept in an enclosure that allows physical interaction (**Fig. 1B, S1A; Table S1**). Because the greatest effect of MDMA at its lowest effective dose (7.5 mg/kg) occurred in the final 10 min of the session (**Fig. 1C, D; Table S1**) we quantified this epoch in all subsequent experiments using a “sociability index” (**See Methods**). While the enhancement of social approach was greatest when both mice received MDMA, a significant enhancement of social preference still occurred when only the free mouse received MDMA (**Fig. 1E; Table S1**). Furthermore, MDMA had equal prosocial effects in both male and female mice (**Fig. S1B; Table S1**) and still caused increased preference for a live mouse compared to a toy mouse placed in the other enclosure (**Fig. S1C; Table S1**). MDMA does not facilitate approach to the social stimulus through an anxiolytic effect, as it generated a mild anxiogenic effect in the elevated-plus maze (**Fig. S1D; Table S1**). Importantly, the lowest dose of MDMA that reliably elicits prosocial behavior (7.5 mg/kg) had minimal locomotor stimulant activity (**Fig. 1F; Table S1**) and did not cause a conditioned placed preference (CPP; **Fig. 1G, H; Table S1**). A higher dose of MDMA (15 mg/kg), however, produced both behaviors (**Fig. 1F, H; Table S1**), which strongly correlate with a drug’s addictive liability (*8, 9*).

**Fig. 1.**
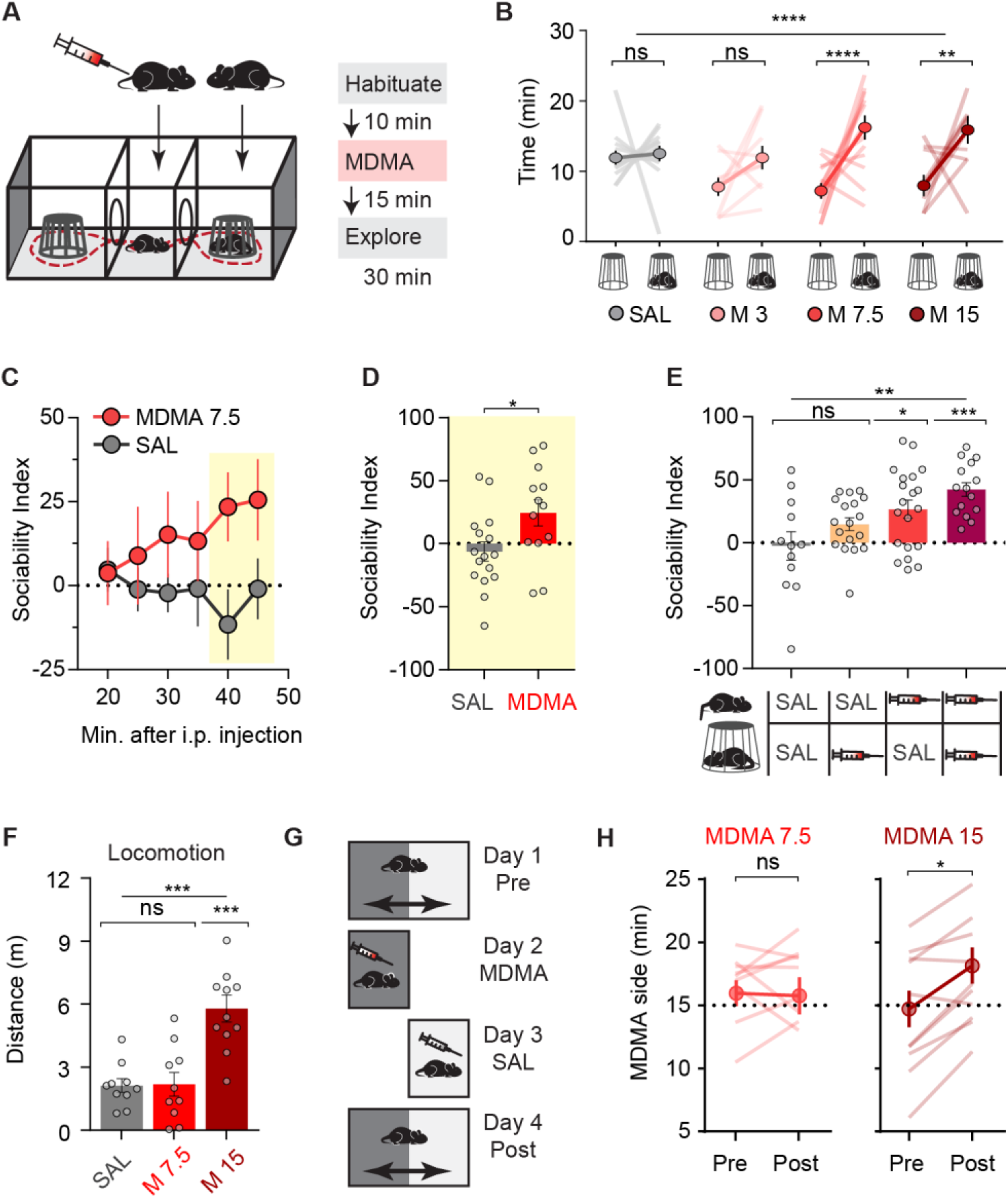
MDMA enhances social preference in mice. (**A**) Three-chamber social interaction assay schematic with experimental timeline. (**B**) MDMA dose-dependently increases time spent with “cup mouse” *versus* “empty cup” (N=9-13). (**C**) Time course of social preference during 30 min exploration after lowest effective dose of MDMA (7.5 mg/kg) *versus* saline (N=14-16). MDMA effect is largest in final 10 min of exploration (yellow box). (**D**) Summary of sociability index in final 10 min. (**E**) MDMA’s prosocial effect as function of mice receiving MDMA or saline injections (N=12-20). (**F**) Lowest effect dose of MDMA in three-chamber assay does not increase locomotor activity, in contrast to higher dose (15 mg/kg; N=10-11). (**G**) Conditioned place preference (CPP) schematic using a single pairing of context with MDMA. (**H**) Preference for MDMA-paired side, pre- and post-conditioning (N=10-11). Low dose MDMA does not induce CPP (*left*), whereas a higher dose does (*right*). Data shown are means ± s.e.m. Significance was determined for each comparison (statistical test): across groups (one-way ANOVA, unmatched) for (B), (E), (F); across group time courses (two-way ANOVA, ordinary) for (C); between groups (unpaired t-test) for (D); within group (paired t-test) for (H); all planned *post hoc* between-group comparison (t-test with Sidak correction for multiple comparisons). *P<0.05; **P<0.01; ***P<0.001; ****P<0.0001; ns, P>0.05.

Several neuromodulatory systems have been implicated in MDMA’s behavioral effects, including 5-HT, DA, and Oxt (*1, 3, 6, 12*), all of which have been suggested to play roles in both social behavior and addiction (*9, 15, 16, 18*). The highest affinity binding of MDMA is to SERT (*6, 7*) leading to supraphysiological 5-HT release through a reverse-transport mechanism (*19–21*). To test if this interaction is required for MDMA’s prosocial effects, we pretreated subjects with the selective serotonin reuptake inhibitor (SSRI) (S)-citalopram (SCIT). SCIT binds to SERT and thereby inhibits MDMA binding but alone does not cause large increases in 5-HT (*20*). While SCIT treatment alone did not alter social preference, it prevented the increase in social approach normally elicited by MDMA (**Fig. 2A; Table S2**).

**Fig. 2.**
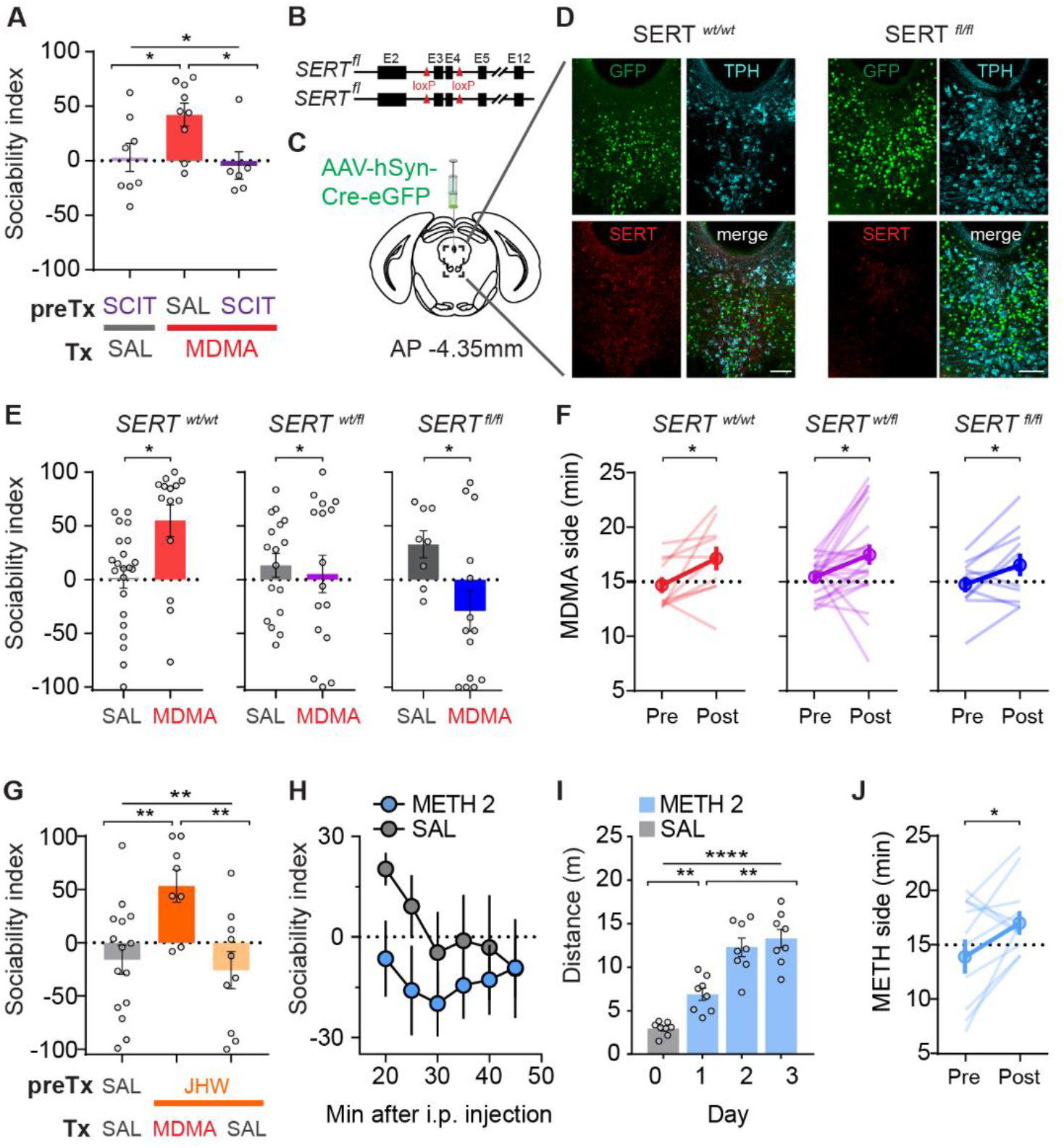
MDMA’s interaction with SERT accounts for its prosocial, but not its rewarding, effects. (**A**) Pre-treatment with the SSRI, SCIT, blocks MDMA’s prosocial effect (N=6-9). (**B**) Schematic of SERT cKO mouse, which has exons 3 and 4 floxed. (**C**) Schematic of AAV containing Cre-eGFP injected into the DR of adult SERT^wt/wt^, SERT^fl/wt^, and SERT^fl/fl^ mice. (**D**) Sample images of immunohistochemistry of Cre-eGFP-injected DR from a SERT^fl/fl^ mouse and wild-type littermate; coronal brain section represents area enclosed by dashed box in **C**. 5-HTergic cells visualized by tryptophan hydroxylase (TPH). (eGFP, green; anti-SERT, red; anti-TPH, cyan; scale bar = 100 μm) (**E**) MDMA (7.5 mg/kg) retained its prosocial effect in Cre-eGFP-injected wild-type mice, but not in heterozygous or homozygous SERT cKO mice (N=8-22). (**F**) High dose MDMA (15 mg/kg) elicits CPP in all three groups of mice from **E** (N=11-22). Preference for MDMA-paired side is shown pre- and post-conditioning. (**G**) MDMA retains its prosocial effect after pre-treatment with a selective DAT inhibitor, JHW-007 (10 mg/kg; N=8-15). (**H**) Time course of sociability index after METH administration (2 mg/kg) *versus* saline (SAL; N=9-10). (**I** and **J**) METH (2 mg/kg) generates locomotor sensitization when given on successive days (**I**; N=8) as well as CPP (**J**; N=12). Data shown are means ± s.e.m. Significance was determined for each comparison (statistical test): across groups (one-way ANOVA, unmatched) for (A), (G); between groups (unpaired t-test) for (E); within group (paired t-test) for (F), (J); across group time courses (two-way ANOVA, ordinary) for (H); within group time course (one-way ANOVA, repeated measures) for (I); all planned *post hoc* between-group comparison (t-test with Sidak correction for multiple comparisons). *P<0.05; **P<0.01; ****P<0.0001; ns, P>0.05.

To independently and more directly test the importance of SERT in mediating MDMA’s prosocial effect, we injected SERT conditional knockout (cKO) mice (SERT^fl/fl^), (**Fig. 2B**) with a Cre recombinase-expressing adeno-associated virus (AAV-Cre) into the dorsal raphe (DR), a major 5-HTergic brain nucleus (**Fig. 2C**). This manipulation dramatically reduced SERT levels in the DR (**Fig. 2D, S2**) and prevented MDMA’s prosocial effect in heterozygous SERT^fl/wt^ mice (**Fig. 2E; Table S2**). Homozygous SERT^fl/fl^ mice that had received DR AAV-Cre injection displayed an increase in baseline social approach following saline injections with no consistent response to MDMA (**Fig. 2E; Table S2**), findings that are consistent with increases in baseline 5-HT levels after SERT KO (*21, 22*). In marked contrast to these findings, genetic deletion of SERT from DR neurons did not influence MDMA-induced CPP in either heterozygous or homozygous SERT cKO mice (**Fig. 2F; Table S2**).

Because MDMA also releases DA via reverse transport due to binding at DAT (*21, 23*), we next examined whether DAT is required for MDMA’s prosocial action. Pretreating subjects with the atypical DAT inhibitor JHW-007, which prevents cocaine’s behavioral effects (*24*), did not influence MDMA’s prosocial effect (**Fig. 2G; Table S2**). Next, we tested methamphetamine (METH), which is structurally related to MDMA and releases DA due to high affinity for DAT, but does not interact with SERT (*7*). METH (2 mg/kg) had no prosocial effect (**Fig. 2H; Table S2**), but did generate both locomotor sensitization (**Fig. 2I; Table S2**) and CPP (**Fig. 2J; Table S2**). These results demonstrate that the MDMA- and METH-elicited behaviors that correlate with addiction (*8, 9*) are largely due to DAT binding and the consequent increase in DA release. In contrast, all results thus far suggest that the prosocial effect of MDMA only requires binding to SERT, not DAT.

Where in the brain does MDMA act to elicit its prosocial effect? Because increasing 5-HT release in the NAc promotes sociability (*16*), we hypothesized that MDMA’s interaction with SERT specifically in the NAc could fully account for MDMA’s prosocial effect. Consistent with this prediction, infusing SCIT bilaterally into the NAc completely blocked the prosocial effect of systemic MDMA (**Fig. 3A, S3A; Table S3**) whereas infusion of SCIT into the VTA, which receives dense DR 5-HT innervation (*25*), had no effect on this behavioral action of MDMA (**Fig. 3B, S3B; Table S3**). Furthermore, direct intra-NAc infusion of MDMA produced a prosocial effect comparable to that elicited by systemic MDMA administration (**Fig. 3C; Table S3**). In contrast, intra-NAc SCIT had no effect on MDMA-induced CPP, which was fully blocked by the dopamine type 2 receptor (D2R) antagonist, raclopride (**Fig. 3D; Table S3**) (*10, 11*). These results suggest that MDMA acting solely on SERT in NAc 5-HT terminals accounts for its prosocial effects.

**Figure 3.**
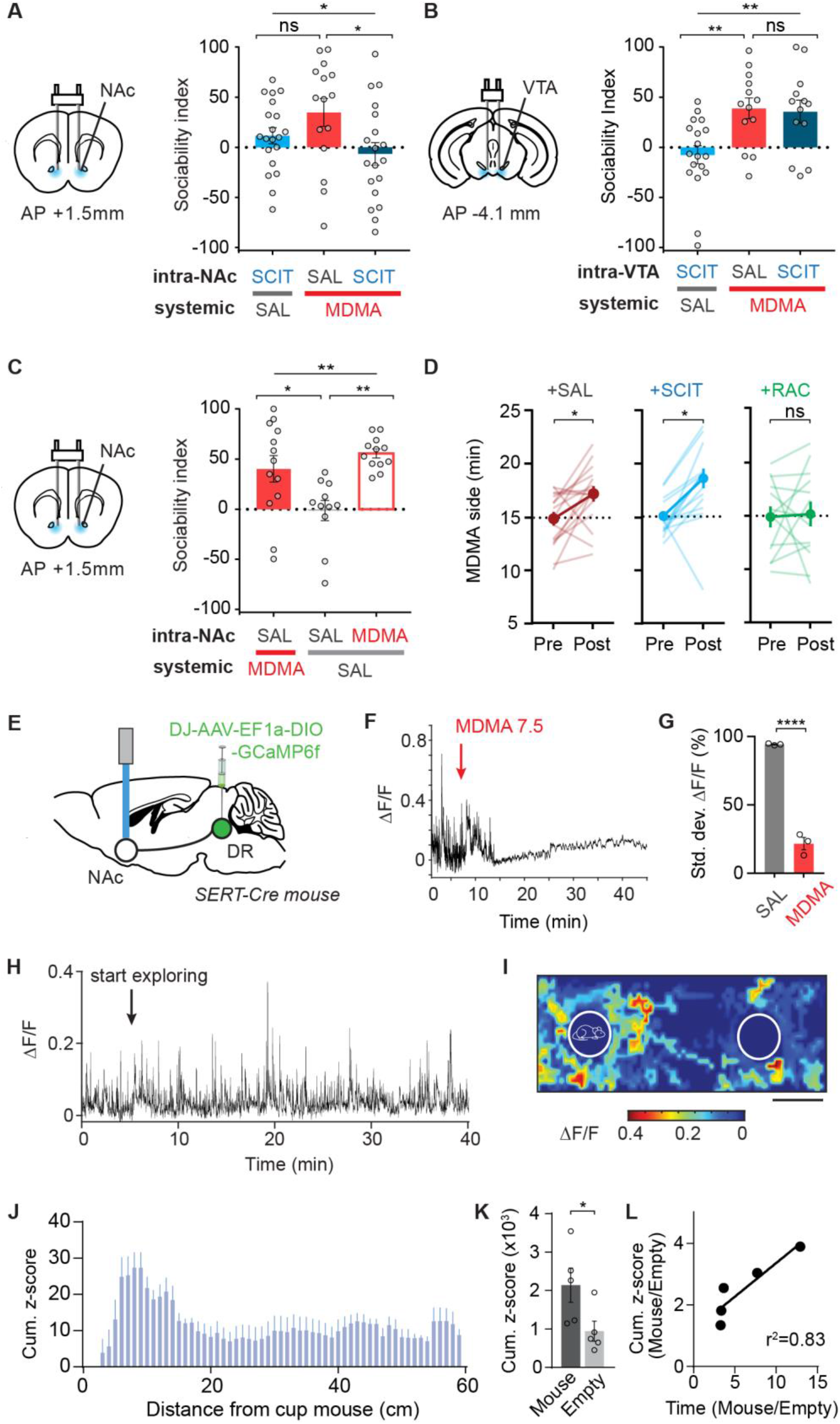
MDMA’s prosocial, but not its rewarding effect, requires interaction with SERT in NAc. (**A**) *Left*, Schematic of drug infusion into the NAc. *Right*, Infusion of SCIT into the NAc blocks MDMA’s prosocial effect (N=15-20). (**B**) *Left*, Schematic of drug infusion into the VTA. *Right*, Infusion of SCIT into the VTA, did not influence MDMA’s prosocial effect (N=13-18). (**C**) *Left*, Schematic as in **A**. *Right*, Infusion of MDMA into the NAc has robust prosocial effect (N=11-13). (**D**) *Left, Center*, Infusion of SAL or SCIT into the NAc does not block CPP elicited by MDMA (15 mg/kg; N=16). *Right*, Infusion of the D2R antagonist, raclopride (RAC), into the NAc blocks MDMA-induced CPP (N=13). (**E**) Schematic of fiber photometry experiments. GCaMP6f was expressed in DR 5-HT neurons to allow imaging of their projections within the NAc during the three-chamber assay. (**F** and **G**) MDMA (7.5 mg/kg) quenches GCaMP6f fluorescence of SERT+ terminals in the NAc. **F**, sample trace. Arrow denotes i.p. MDMA injection. **G**, Summary graph of standard deviation (SD) of ΔF/F during 5 min epoch after either saline (SAL) or MDMA (7.5 mg/kg), normalized to SD of ΔF/F during the 5 min before injection (N=3). (**H**) Sample trace illustrating GCaMP6f transients from SERT+ terminals in the NAc during three-chamber exploration. (**I**) Spatial heat map example. The maximal ΔF/F occurring for each explored area of the three-chamber apparatus is shown for one 30 min session. Outlines of mouse cup (left) and empty cup (right) are drawn in. (**J**) Summary graph of z-scored fluorescence as a function of distance from the mouse cup. (**K**) Summary graph of cumulative z-scored fluorescence for the area around the mouse cup *versus* empty cup (N=5 mice). (**L**) Ratio of mouse cup to empty cup-related GcaMP6f fluorescence as function of the ratio of time spent in the same two areas. Data shown are means ± s.e.m. Significance was determined for each comparison (statistical test): across groups (one-way ANOVA, unmatched) for (A), (B), (C); within group (paired t-test) for (D); between groups (unpaired t-test) for (G), (K); univariate correlation (linear regression) for (L); all planned posthoc between-group comparison (t-test with Sidak correction for multiple comparisons). *P<0.05; **P<0.01; ****P<0.0001; ns, P>0.05.

To further test this hypothesis, we expressed the fluorescent calcium indicator, GCaMP6f, in DR 5-HT neurons and recorded fluorescence changes in 5-HT fibers in the NAc during administration of MDMA (**Fig. 3E-G; S3C**). MDMA binding to SERT causes reverse transport of 5-HT and collapse of the normal pH gradient between 5-HT synaptic vesicles and the cytoplasm (*19*). Because GCaMP6f fluorescence is pH sensitive (*26*), by acidifying terminals upon which it acts (*19*), MDMA should quench this fluorescence. Consistent with this prediction, systemic MDMA administration caused a rapid, large reduction of GCaMP6f fluorescence in DR 5-HT inputs in NAc (**Fig. 3F, G; Table S3**). In the absence of MDMA, activity in NAc 5-HT inputs increased when the free test mouse approached the “cup” mouse in a manner that inversely correlated with their distance apart (**Fig. 3H-K; Table S3**). In addition, the magnitude of this increase correlated with the degree of social preference exhibited by the test mouse (**Fig. 3L; Table S3**). These findings provide further evidence that activity in DR 5-HT inputs in the NAc encodes some component of social preference and MDMA acts directly on these inputs.

MDMA’s effects on social interactions have been suggested to involve Oxt (*3*), a neuropeptide that plays important roles in a variety of social behaviors (*18*). However, pretreating mice with the Oxt receptor (OxtR) antagonist L-368,899 at a dose (5 mg/kg) that blocks Oxt-dependent behaviors (*15*), had no effect on MDMA-induced sociability (**Fig. 4A: Table S4**). Higher, repeated systemic doses of L-368,899 or intra-NAc infusion of either L-368,899 or the more selective and potent OxtR antagonist, L-371,257, also had no influence on MDMA’s prosocial effect (**Fig. 4A, B; Table S4**). Finally, MDMA still produced its normal prosocial effect in mice in which OxtRs were genetically deleted in 5-HT neurons by crossing the OxtR^fl/fl^ mouse line with the SERT-Cre mouse line (**Fig. 4C, Table S4**). These findings provide strong evidence that OxtRs are not required for the prosocial effect of MDMA and are consistent with previous findings that Oxt release in the NAc promotes social reward by increasing 5-HT release (*15*).

**Fig. 4.**
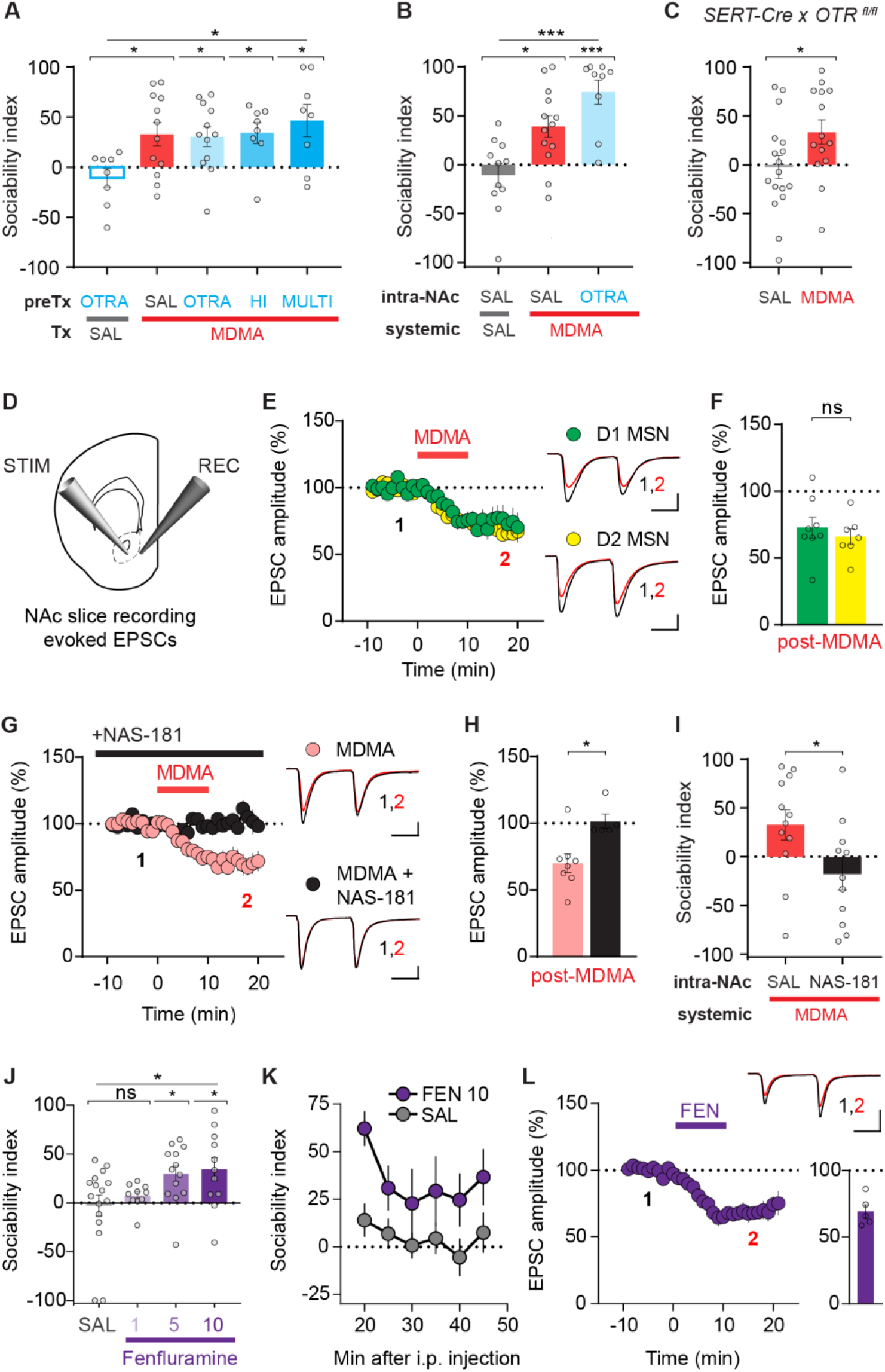
MDMA’s prosocial effect requires 5-HT release and activation of 5-HT1b receptors in NAc. (**A**) Several dose regimens of the OxtR antagonist (OTRA), L-368,899, did not influence MDMA’s prosocial effect: OTRA (5 mg/kg once); HI (10 mg/kg once); MULTI (5 mg/kg every 12 h for 48 h preceding MDMA). All groups differed significantly from OTRA given alone (N=8-12). (**B**) Direct infusion of OTRA into the NAc did not influence MDMA’s prosocial effect. Results with two OTRAs, L-368,899 and L-371,257, are pooled (N=9-13). (**C**) MDMA retained its prosocial effect in mice with selective deletion of OxtRs in SERT+ cells (N=14-17). (**D**) Schematic for slice electrophysiology from coronal brain slices containing NAc. (**E**) *Left*, Summary graph of effects of bath application of MDMA (10 μM) on EPSCs recorded from D1 and D2 MSNs. *Right*, Sample EPSC traces taken from time points (1) and (2) as shown. (**F**) Summary graph shows MDMA depressed EPSCs in D1 and D2 MSNs to a similar extent (N=7-8 cells, 6-7 mice). (**G**) Summary graphs showing the 5-HT1b receptor antagonist, NAS-181 (20 μM), blocks MDMA’s effect on NAc EPSCs. Pooled data from D1 and D2 MSNs are shown. *Right*, sample traces, as in **E**. (**H**) Summary graph of data in **G** (N=5-8 cells, 3-5 mice). (**I**) MDMA’s prosocial effect is blocked by NAS-181 infused into the NAc (N=11-12). (**J**) The 5-HT-releasing drug fenluramine (1,5, 10 mg/kg) has a dose-dependent prosocial effect (N=10-16). (**K**) Time course of sociability index after fenfluramine (FEN) injection (10 mg/kg; N=11) *versus* saline (SAL; N=16). (**L**) Effects of bath application of FEN (10 μM) on EPSCs recorded from NAc MSNs (N=5, 3 mice). *Top right*, Sample EPSC traces taken from time points (1) and (2) as shown. *Bottom right*, Summary graph of FEN-mediated depression of EPSCs. Scale bars: 100 pA, 25 ms in E, G; 200 pA, 25 ms in L. Data shown are means ± s.e.m. Significance was determined for each comparison (statistical test): across groups (one-way ANOVA, unmatched) for (A), (B), (J); between groups (unpaired t-test) for (C), (F), (G), (I); across group time courses (two-way ANOVA, ordinary) for (K); all planned posthoc between-group comparison (t-test with Sidak correction for multiple comparisons). *P<0.05; ***P<0.001; ****P<0.0001; ns, P>0.05.

To determine if MDMA’s effect on NAc physiology mimics that of 5-HT (*15, 27*), as predicted by our hypothesis, we made whole cell voltage clamp recordings from visually identified NAc medial shell D1R and D2R expressing medium spiny neurons (MSNs) in acute brain slices and bath applied MDMA while stimulating excitatory inputs (**Fig. 4D**). MDMA (10 μM) precisely mimicked the actions of 5-HT in that it elicited a long-term depression (LTD) of excitatory postsynaptic currents (EPSCs) in both D1- and D2-MSNs (**Fig. 4E, F; Table S4**), which was prevented by prior application of the 5-HT1b receptor antagonist, NAS-181 (**Fig. 4G, H; Table S4**). Intra-NAc infusion of NAS-181 *in vivo* also blocked MDMA’s prosocial effect (**Fig. 4I; Table S4**), indicating that the behavioral and electrophysiological effects of MDMA-evoked 5-HT release requires activation of NAc 5-HT1b receptors.

A role for 5-HT release and specific 5-HT receptor subtypes in mediating MDMA’s prosocial effect has previously been suggested (*3*). However, this work did not address the locus of MDMA’s action nor did it define if the mechanisms mediating MDMA’s acute reinforcing effect are the same as those mediating its effects on sociability. Our demonstration that MDMA’s addictive liability appears to be due entirely to its DA releasing properties predicts that a drug, which mechanistically functions like MDMA but binds only to SERT and not DAT, should have the same prosocial effect with no acute reinforcing properties. D-fenfluramine (FEN) exhibits these properties (*7*), has been approved for investigational use in humans (*28*) and, importantly, has no acute reinforcing effect when tested in multiple species (*7, 29*). We therefore examined FEN and found that, like MDMA, it enhanced social preference dose-dependently (**Fig. 4J, K; Table S4**), and generated an identical electrophysiological signature (**Fig. 4L; Table S4**).

The importance of our conclusions for understanding the mechanisms mediating MDMA’s unique prosocial actions in humans depends on the assumption that the neural mechanisms we have elucidated in mice reflect those mediating MDMA’s actions in the human brain. Several lines of evidence support this assumption. First, the consistent, robust effects of MDMA in the three chamber assay parallels the prosocial effect that is central to the human experience. Second, at higher doses, MDMA’s psychomotor stimulant and acute reinforcing actions predict addictive liability (*8, 9*), which has been documented in human MDMA abusers (*1*). Third, human pharmacology experiments suggest that aspects of MDMA’s subjective effects are sensitive to both an SSRI and D2R antagonist (*12*), findings that mimic our more precise manipulations in mice. Fourth, the prosocial effects of Oxt administration in humans are qualitatively different from those of MDMA (*3*), consistent with the lack of a role for Oxt in mediating MDMA’s acute prosocial effects in our assays. Finally, despite the toxicity associated with long term use (*7*), early trials showed promise using FEN to treat social deficits in children with autism (*30*).

Given MDMA’s long history of abuse and potential toxicity, should it prove efficacious in treatment of neuropsychiatric disorders such as PTSD and autism spectrum disorders, it would be prudent to develop drugs or other therapies that mimic its prosocial effects with reduced potential associated morbidities. While much remains unknown about the detailed neural mechanisms by which increased 5-HT release in the NAc promotes sociability, our findings provide a defined mechanistic basis for the further development and testing of agents that positively influence social interactions in a therapeutic manner.

## Acknowledgments

We thank Dr. Jai Pollepali, Dr. Kevin Beier, Ms. Marija Pavlovic, Dr. Matthew A. Wright and Dr. Roberto de Gregorio for technical assistance.

## Author contributions

B.D.H. and R.C.M. initiated the project and designed the experiments; B.D.H., J.S.S., M.D.T., D.F.C.P., E.E.S. and J.J.W. performed and analyzed behavioral assays; B.D.H. performed and analyzed fiber photometry assays; B.D.H. and P.H. performed and analyzed electrophysiological assays. B.D.H. and R.C.M. wrote the paper which was reviewed and edited by all authors.

## Data and materials availability

All data are reported in the main text and supplementary materials. All data, code and materials are stored at Stanford and available upon reasonable request.

