## Supplementary materials for "Distinct neural mechanisms for the prosocial and rewarding properties of MDMA"

**This PDF file includes:**

Materials and Methods
Fig S1 - S3
Tables S1 – S4

**Materials and Methods**

**Animals.** Male and female C57Bl/6 mice, aged 8-16 weeks, (Jackson Laboratory; stock 000664), and the following transgenic lines were used:

- $Slc6a4^{fl/fl}$ (homozygous), $Slc6a4^{fl/wt}$ (heterozygous), and wild-type littermate mice, male and female ($SERT^{fl/fl}$, $SERT^{fl/wt}$, $SERT^{wt/wt}$, gift from Ji-Ying Sze, Albert Einstein College of Medicine) (*31*);
- Tg(Slc6a4-cre)ET33Gsat male mice (SERT-Cre, GENSAT Project at Rockefeller University; MGI: 3836639) (*32*);
- B6.129(Cg)-$Slc6a4^{tm1Kpl}$/J male mice (SERT-KO, Jackson Laboratory Stock #: 008355)
- $Oxtr^{tm1.1Wsy}$/J homozygous ($OxtR^{fl/fl}$, Jackson Laboratory stock #:008471; MGI:3800791) (*33*);
- SERT-Cre x $OTR^{fl/fl}$ male mice were generated in the Malenka lab using a breeding strategy previously described for producing DAT-Cre x OTR $^{fl/fl}$ mice (*34*).
- Drd1a–td Tomato / Drd2– eGFP BAC double transgenic mice were backcrossed to wild-type C57Bl/6 mice (Jackson Laboratory, as above) and offspring (*35*) were used for electrophysiology experiments.

All mice were maintained on a C57Bl/6 background, and housed on a 12-hour light/dark cycle with food and water *ad libitum*. All behavioral experiments were conducted during the same circadian period (7:00am to 4pm). All procedures complied with animal care standards set forth by the National Institute of Health and were approved by Stanford University's Administrative Panel on Laboratory Animal Care and Administrative Panel of Biosafety. Experimental group sizes were determined by power analyses using the effect size observed during initial exploratory experiments, taking into account historical accuracy within our group for stereotactic targeting of a given brain region.

**Stereotactic surgery.** All surgeries were performed in a sterile, temperature-controlled environment. Mice were anesthetized with inhaled isoflurane (1-1.5%), maintaining spontaneous respiration. Mice were positioned in a stereotactic head frame (Kopf Instruments, Tujunga, CA; Model 940), and the skull surface was exposed. Craniotomy for viruses and implants was made with a 0.5 mm drill bit (Fine Science Tools, Foster City, CA; 19007-05). Implants were stabilized with the insertion of 1 to 2 skull screws (Antrin Miniature Specialties; 00-90 x 1/16), and sequential application of C&B Metabond® (Parkell, Edgewood, NY) and Dual Cure Resin Ionomer (DenMat, USA; Geristore®, no. 4506). Local anesthetic (lidocaine 0.5%; APP Pharmaceuticals, LLC) was infiltrated into the scalp incision and meloxicam (Boehringer Ingelheim Vetmedica, Inc.; 5 mg/kg s.c.) was given for postoperative analgesia.

**Virus injection.** A syringe (Hamilton, Reno, Nevada; Model 85 RN SYR) with a thirty-three gauge needle (VWR, Radnor, PA; #7762) was used to infuse 400 nL of virus into the dorsal raphe (DR), bregma coordinates: anteroposterior (AP) -4.36 mm, mediolateral (ML) 0.00 mm, dorsoventral (DV) -3.00 mm. Virus was infused at a rate of 100 nL per minute, and the injection needle was withdrawn 5 minutes after the end of the infusion. Viruses used in this study were purchased from the Stanford Neuroscience Gene Vector and Virus Core and include DJ-AAV-hSyn-Cre-eGFP and DJ-AAV-EF1a-DIO-GCaMP6f. For behavioral experiments involving virus injection, behavior was assessed at least 4 weeks after surgery.

**Drug infusion cannulae implants.** Double lumen, twenty-six gauge threaded cannula guides for bilateral drug infusion were custom ordered (PlasticsOne, Roanoke, VA). Cannula guides were lowered into the craniotomy sites to a position 1.5 mm above the target structure. After securing the cannula guide (see above), a bilateral stylet ('dummy cannula') was inserted into the infusion ports and the device was sealed with a threaded aluminum cap. Cannula guide dimensions were: (nucleus accumbens [NAc] infusions) 1.8 mm separation, 8 mm pedestal, cut 3.5 mm below pedestal; (ventral tegmental area [VTA] infusion) 1.0 mm separation, 8 mm pedestal, cut 3.8 mm below pedestal. Bregma coordinates for cannula guide implantation were: (NAc) AP +1.50 mm, ML ± 0.90 mm, DV -2.80 mm; (VTA) AP -3.05 mm, ML ± 0.50 mm, DV -3.13 mm. For drug infusion studies, behavioral experiments were performed 2-4 weeks after surgery.

**Optical fiber implants.** Fiber optic implants (Doric Lenses, Quebec, Canada; 400 μm thick, 0.48 NA, custom cut to length) for fiber photometry experiments in the NAc were lowered into the craniotomy site and secured, as described above, at bregma coordinates: AP -1.50 mm, ML -0.90mm, DV -4.3 mm.

**Behavioral testing randomization and blinding procedures.** For all behavioral tests (three-chamber social approach, locomotion, conditioned place preference, elevated plus maze) experimental groups were interleaved with control groups. All behavioral experiments were conducted in a blind manner such that: animals were randomized by cage prior to surgery and experiments; for experiments involving transgenic animals, the experimenter was blind to the genotype; performance of a behavioral experiment and analysis of the resulting data were performed by separate individuals; and analysis was performed without knowledge of experimental group assignment. For three-chamber social approach experiments, mice were typically used once in an experimental group and once in a control group, with treatments occurring at least one week apart to allow for drug washout. We verified that MDMA (7.5 mg/kg) could elicit a similar prosocial effect when given one week apart (data not shown).

**Three-chamber social approach.** The three-chamber apparatus was constructed in the lab using .635 cm thick sheets of clear extruded acrylic for walls, and Komatex® for floors (white) and barrier walls (black; TAP Plastics, Mountain View, CA). The three-chamber apparatus (L x W x H; 72 cm x 23 cm x 25 cm) consisted of two outer chambers (28 cm x 23 cm) connected by a center chamber (16 cm x 23 cm). Translucent acrylic walls with 6.5 cm diameter holes for mouse passage, drilled 1 cm from the chamber floor, defined the center chamber. These walls were omitted from the fiber photometry experiments to allow for patch cord connection to the free mouse. 'Cup mice' were kept in one of the two compartments under a 10 cm diameter inverted metal pencil cup.

All mice were handled daily for three consecutive days, habituated to the behavioral testing room, and habituated to i.p. saline injection prior to behavioral testing. Social approach testing was performed similar to previously described methods (*17*). On the test day, experimental mice ('free') were habituated to the 3 chamber apparatus for 10 minutes, with an empty inverted wire pencil cup placed in each outer chamber. Mice were briefly returned to their home cage, and after receiving test drug or saline i.p. injections, or an intracerebral infusion, they were replaced into the 3 chamber apparatus, now confined to the center chamber by removable opaque barrier walls. A conspecific, age and sex-matched, wild-type, unfamiliar (non-cagemate) mouse was placed under one of the inverted cups ('cup mouse'). The 'cup mouse', except in the experiment described in **Fig. 1E**, received no drug treatment. Fifteen minutes after i.p. injection, dividing walls were removed and ‘free’ mice were allowed to explore the entire apparatus for thirty minutes. At the end of the session, mice were returned to their home cage. Between experimental sessions, cups and the 3 chamber apparatus were sprayed with Virkon-S® and 70% ethanol, and wiped down. The ‘cup mouse’ side of the apparatus and the cups used to hold mice were regularly rotated to minimize the effect of residual scent cues on behavioral testing.

Video was acquired by a PC-controlled ceiling mounted digital camera, and analyzed offline using Biobserve tracking software (Biobserve GmbH, Bonn, Germany). Fidelity of automated mouse tracking was verified for each experiment, and time spent, per minute, was quantified for each of the three chambers. To enable comparisons across treatment groups and minimize the impact of variable exploration times between mice, we calculated a ‘sociability index’ for each mouse as 100 x (‘cup mouse’ time – ‘cup’ time)/ (‘cup mouse’ time + ‘cup’ time). Mice were excluded from further analysis if they either failed to explore both ‘cup’ and ‘cup mouse’ chambers, or if they spent more than 75% of allotted time in the center chamber, not exploring either cup.

**Locomotor assay.** The locomotor testing chamber (Med Associates, Fairfax, VT) was an acryclic box, internal dimensions of 28 cm x 28 cm x 20 cm. Mice were allowed to explore for 60 minutes, starting immediately after i.p. injection. Mouse movement was tracked by an array of infrared beam break counters, recorded using Med Associates software, and analyzed offline.

**Conditioned Place Preference (CPP).** Two equal-sized compartments were created within each locomotor testing chamber described above. Each compartment had a distinct floor, either clear, textured acrylic (‘A’ side) or smooth, black acrylic (‘B’ side). Left/right positioning of the floors was alternated between conditioning chambers. Conditioning experiments involving MDMA took place over 4 consecutive days.

Day 1, Pre-test: for 30 minutes, the mouse had free access to both compartments, separated by a clear acrylic divider with a 7 cm diameter hole for passage. Baseline preference was measured as the amount of time spent per side. Mice were pseudo-randomly allocated to either the ‘A’ or ‘B’ side for drug conditioning such that: net baseline preference within an experimental group was as close to 15.0 minutes as possible; equal numbers of mice were assigned to ‘A’ and ‘B’ sides.

Day 2, 1st conditioning day: for 60 minutes, the mouse was sequestered to either ‘A’ or ‘B’ side by a solid transparent divider, starting 15 minutes after receiving test drug or saline treatment. For experiments involving intracerebral infusions. Any mouse receiving test drug by intracerebral infusion on one conditioning day received intracerebral infusion of saline on the other conditioning day. Half the mice within an experimental group received test drug on Day 2, the other half on Day 3.

Day 3, 2nd conditioning day: as on Day 2, with treatment (test drug or saline) and compartment (‘A’ or ‘B’) switched.

Day 4, Post-test: as on Day 1.

For methamphetamine CPP, two rounds of conditioning were performed, for a total of 6 consecutive days.

**Elevated Plus Maze.** The elevated plus maze consisted of two metal rails (dimensions, 80 cm x 5 cm) perpendicular to each other and intersecting at their centers, as show in **Fig. S2D**, raised 50 cm above ground level. Opaque black walls (15 cm high) surrounded two arms, while the other arms were open. MDMA (7.5 mg/kg) was given 25 minutes prior to a 10-minute exploration period. Video was acquired and analyzed as described for the three-chamber assay. Time spent in each arm was quantified and net time in ‘closed’ and ‘open’ arms was compared.

**Drug treatments for behavioral experiments.** Drugs were administered i.p. at a volume of 0.01 mL/g. MDMA (3-15 mg/kg; Sigma), (S)-citalopram (7 mg/kg;’escitalopram oxalate’, Tocris), methamphetamine (2 mg/kg; Sigma), L-368,899 (5-10 mg/kg; Tocris) and d-fenfluramine (1-10 mg/kg; Tocris) were all dissolved in 0.9% normal saline. JHW-007 (10 mg/kg; Tocris) was dissolved in 2.5% DMSO and 0.9% normal saline. For experiments involving two drugs given i.p., pre-treatments ([S]-citalopram, L-368,899, JHW-007) were uniformly given 10 minutes prior to MDMA i.p. injection.

For intracerebral infusions, all drugs except L-371,257 were dissolved in 0.9% normal saline to achieve a concentration of (S)-citalopram (0.5 µg), MDMA (0.5 µg), (S[-]-raclopride (+)-tartrate (0.5 µg; Sigma), L-368,899 (0.5 µg), or NAS-181 (0.5 µg; Sigma) in 500 nL infusate, delivered bilaterally at a rate of 350 nL per minute. L-371,257 (0.25 µg; Tocris), was dissolved in 5% DMSO and 0.9% normal saline and infused in the same manner as other drugs. This vehicle solution was infused into the NAc alone or in combination with either saline or MDMA (7.5 mg/kg), for a portion of the experiments represented in **Fig. 4B**.

For drug infusion, the dummy cannula was replaced with a bilateral infusion cannula measuring 1.5 mm longer than its corresponding cannula guide. Infusions were delivered by inserting drug-primed bilateral infusers into the implanted cannula guide. Infusers were connected via PE-50 tubing to a 23 gauge needle of a syringe (Hamilton, Reno, Nevada; Model 1701 RN and 7786-01) mounted in a dual syringe pump (Harvard Apparatus, Holliston, MA; Model 55-2222). After a 90-sec infusion, the infuser was held in place for an additional 30 seconds, then the 'dummy cannula' was re-inserted into the cannula guide and the implant was re-sealed with an aluminum cap. For experiments involving both i.p. injection and intracerebral infusion, i.p. injection was done just after the infusion.

**Fiber Photometry.** AAV-DJ-DIO-GCaMP6f was infused into the DR and a fiber optic implant was advanced and secured in the NAc using the same stereotactic coordinates noted above. After allowing 3-4 weeks for viral expression, mice were first habituated to the fiber photometry apparatus including attachment of a 2 m patch cord for 30 minutes, and then tested on a subsequent day. Behavior testing consisted of the three-chamber assay, as described above, with continuous video and fiber photometry acquisition. Fiber photometry data was acquired with Synapse software controlling an RZ5P lock-in amplifier (Tucker-Davis Technologies). GCaMP6f excitation was achieved with frequency-modulated 473nm and 405nm LEDs (Doric), to stimulate $Ca^{2+}$-dependent and isosbestic emission respectively. All optical signals were band-pass filtered with a Fluorescence MiniCube FMC4 (Doric), emission was measured with a femtowatt photoreceiver (2151; Newport), and signal was digitized at 6 kHz.

Signal processing was performed with Matlab (Mathworks, Inc.). To correct for motion artifact and fluorescent bleaching, signals generated from the two excitation wavelengths ($F_{473}$ and $F_{405}$) were used to calculate $\Delta F/F = (F_{472} - F_{405}) / \text{mean}(F_{405})$. For instances where fluorescent decay of $F_{473}$ and $F_{405}$ appeared to have different time constants (i.e. $F_{473}/F_{405}$ was markedly nonlinear), each signal $F_{\lambda}$ was debleached by fitting with a mono- or bi-exponential decay function; $\Delta F/F_{\lambda}$ was calculated for each wavelength as $F_{\lambda}/\text{mean}(F_{\lambda})$; the final corrected $\Delta F/F$ was then calculated as $\Delta F/F_{472}$ - $\Delta F/F_{405}$. For experiments involving quenching of fluorescence by MDMA, corrections were applied using data preceding drug application. A zerophase digital filter was applied to the resulting signal and data were z-scored for some comparisons as indicated in main text.

Video for fiber photometry experiments in the three-chamber apparatus was acquired as described above. High resolution motion tracking was done with TopScan software (CleverSys, Reston, VA). Z-score or ΔF/F *vs* time, and mouse position *vs* time, were combined to create a map of the maximal z-score observed for each position visited by the test mouse within the three-chamber apparatus over a 30 minute session. For quantitation of fluorescent signal associated with the 'mouse cup' and 'empty cup', maximal z-scores were summed for all positions in a 9 cm x 9 cm area centered on the cup. For the illustrated example (**Fig. 3I**), the maximal z-score map was linearly interpolated by a factor of 10 for presentation.

**Electrophysiology.** Whole cell recordings were performed in 2-4 month old male mice, either Drd1a–td Tomato / Drd2– eGFP BAC double transgenic mice, or wild-type C57Bl/6. Coronal slices containing the NAc, identified by the corpus callosum and anterior commissure, were prepared as previously described (*15*). Briefly, mice were deeply anesthetized with pentobarbital until unresponsive to tail pinch, but still breathing. 95% $O_2$ was supplied by face mask until the left ventricle was cannulated, and the mouse was perfused through this cannula with ice-cold 'sucrose solution' containing (in mM): 215 Sucrose, 2.5 KCl, 20 glucose, 26 $NaHCO_3$, 1.6 $NaH_2PO_4$, 1 $CaCl_2$, 4 $MgSO_4$ and 4 $MgCl_2$. After brain removal, blocked hemispheres were mounted and 250 µm thick slices were cut on a vibratome (Leica, VT1200S) in ice-cold 'sucrose solution'. Slices were immediately transferred to a bath containing artificial cerebrospinal fluid (ACSF), warmed to 35°C, containing (in mM): 124 NaCl, 2.5 KCl, 10 glucose, 26 $NaHCO_3$, 1 $NaH_2PO_4$, 2.5 $CaCl_2$ and 1.3 $MgSO_4$. After 30 minutes, slices were maintained at room temperature for at least one hour prior to transfer to the recording chamber. Cutting and recording solutions were saturated with 95% $O_2$ and 5% $CO_2$, pH 7.4. Experiments were performed at 30.0 ± 0.3 °C.

Upon transfer to the recording chamber, slices were completely submerged and continuously superfused at a flow rate of 3 mL per minute with ACSF containing the $GABA_A$ receptor antagonist, picrotoxin (50 µM, Sigma). A stainless steel monopolar stimulating electrode (2-5 Mohm impedance; FHC, Brunswick, ME) was advanced into tissue, placed approximately 150-300 µm away from the recording site and near the anterior commissure.

Capillary glass pipettes (3-4 Mohm) were filled with an internal solution containing (in mM): 130 $CsMeSO_3$, 10 HEPES, 0.4 EGTA, 5 TEA-Cl, 7 $Na_2$phosphocreatine, 4 MgATP, 0.4 NaGTP, 0.1 Spermine, 4 QX-314 Cl (pH 7.3; 290-295 mOsm). Neurons were patched under direct visualization using IR-filtered differential interference contrast microscopy. Fluorophore-labeled neurons were identified prior to patching using epifluorescence microscopy. Whole-cell, voltage-clamp recordings were performed with a MultiClamp 700A amplifier (Axon Instruments Inc., Union City, CA) whose output signals were filtered at 3 kHz; recordings were digitized at 20 kHz. After establishing intracellular access, cells were held at -70 mV. Series resistance (typically 8-20 MΩ) and input resistance (typically 100-300 MΩ) were continuously monitored throughout each experiment with a -5 mV, 80 ms command pulse delivered prior to each electrical stimulus pair. Cells with more than 20% change in series resistance were excluded from analysis. Series resistance, input resistance, holding current and EPSC amplitude were analyzed on-line using customized software for IgorPro (Wavemetrics Inc., Lake Oswego, OR).

To evoke excitatory postsynaptic currents (EPSCs), square pulse stimuli (200 µs pulse width, 50 msec interval) were delivered via a constant-voltage stimulator every 15 sec. Stimulation intensity was adjusted to evoke EPSCs with amplitudes of 200-500 pA. For each experiment, a baseline (minimum of 10 minutes) was first obtained in which EPSC amplitude did not vary from beginning to end of baseline period by more than 15%. Test drugs were added from stock solutions, maintained on ice, to the ACSF reservoir at a dilution of 1:1000 to achieve the following concentrations: 10 µM MDMA, 20 µM NAS-181, 10 µM d-fenfluramine. MDMA and d-fenfluramine were applied for 10 minutes and then washed out with the ACSF recording solution. NAS-181 was continuously superfused throughout experiments. To quantify the change in EPSC after drug application, the average EPSC amplitude from 15 to 20 minutes post drug application was calculated. Representative traces were generated by average waveforms from 5 consecutive minutes of baseline recordings (from 5 minutes prior to drug application) or from the 15 to 20 minutes post-drug epoch. For presentation, stimulus artifacts were manually removed.

**Immunohistochemistry.** The animals were terminally anesthetized with pentobarbital, transcardially perfused with 10% formalin, and post-fixed overnight in the same solution. The following day, coronal brain sections (60 µM) were cut on a vibratome in 1M phosphate-buffered saline (PBS) solution. The free-floating sections were permeabilized with PBS-0.3%

Triton-X100 for 30 minutes and incubated for 1 hour at room temperature (RT) in a blocking solution with 5% Goat Serum in PBS-1% Triton-X100, and then incubated overnight at 4$^{o}$C with the following primary antibodies (dilution; vendor), in combinations indicated in the text: chicken anti-GFP (1:2000; Aves Labs, USA), rabbit anti-5HT transporter (1:1000; Millipore, USA), and goat anti-TPH-2 (1:1000; Novus, USA). The following day, sections were rinsed in PBS three times for 10 minutes at RT and incubated in the same blocking solution for 1.5 hour at RT with matched secondary antibodies (1:500; ThermoFisher, USA): Alexa Fluor 488 anti-chicken, Alexa Fluor 568 anti-rabbit, and Alexa Fluor 647 anti-goat. The sections were then rinsed three times for 5 minutes and mounted on slides with DAPI Fluoromount-G (SouthernBiotech). Images were acquired on a Nikon confocal microscope at 10x and 40x magnification. Each image was acquired using identical pinhole, gain and laser settings.

**Statistics.** Student's two-tailed t-tests were used to compare two groups. Paired comparisons were performed for CPP assays (pre- *vs* post-conditioning) and real time place preference assays (initial *vs* reversed light-paired compartment); unpaired comparisons were used elsewhere. One-way ANOVA, unmatched, using the Sidak correction for predetermined *post hoc* subgroup comparisons were used when multiple conditions were compared. Two-way ordinary ANOVA was performed for comparisons of multiple treatment conditions over multiple time points. Statistical analyses were performed using Prism 7.04 (GraphPad Software). All data were tested and shown to exhibit normality and equal variances. All data are expressed as mean ± s.e.m.

#### Figure S1

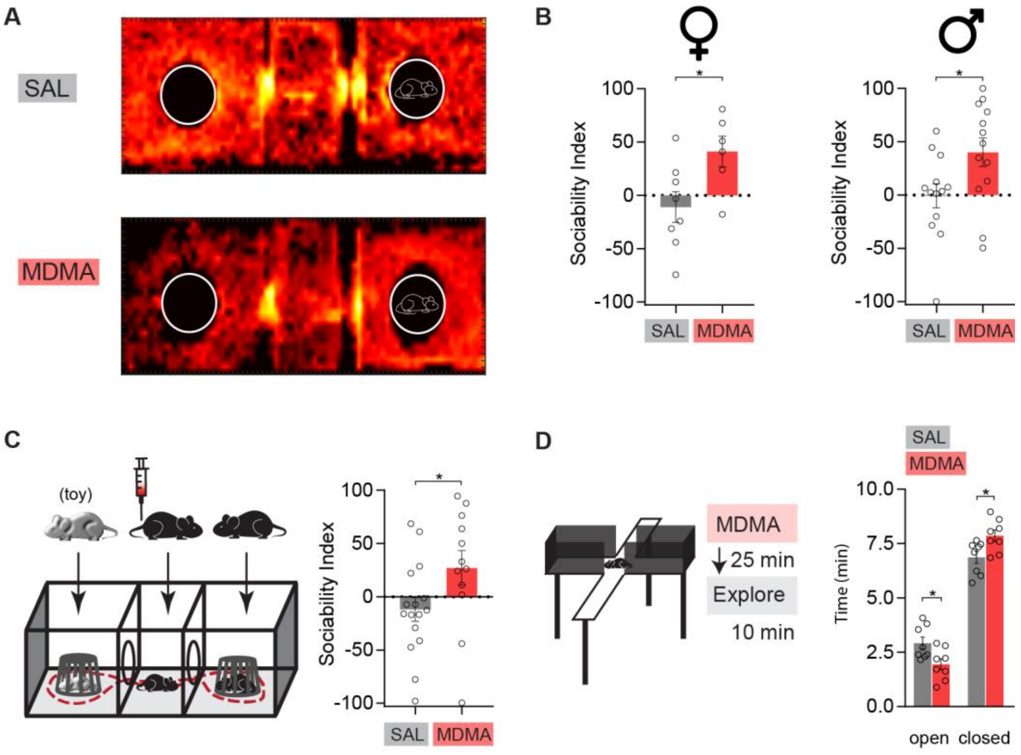

**Fig. S1. Supplementary information supporting data shown in Fig. 1.** (**A**) Sample heat map illustrating the relative amounts of time which a saline- or MDMA-treated mouse spent within each position of the three-chamber testing apparatus over the 30-minute exploration period. Heat map construction: the sum total of visits to each square centimeter was calculated; for visualization, the map (M) was transformed by 1 + log(M); map was interpolated by a factor of 10; a gaussian smoothing function was applied. (**B**) MDMA (7.5 mg/kg) produces a prosocial effect in adult female mice (*left*; N=6-8), comparable to the effect observed in adult male mice (*right*; N=13). (**C**) MDMA (7.5 mg/kg) still produces prosocial effect when an unfamiliar toy mouse (schematic at *left*) is placed in the empty cup (summary graph at *right*; N=12-16). (**D**) *Left*, Schematic of elevated-plus maze and experimental time line. *Right*, Mice treated with MDMA (7.5 mg/kg) spent significantly more time in the closed-arm areas compared to SAL treated mice, suggesting an anxiogenic drug effect (N=8). Data shown are means ± s.e.m. Significance was determined between groups using an unpaired t-test for (B), (C), (D). *P<0.05.

### Figure S2

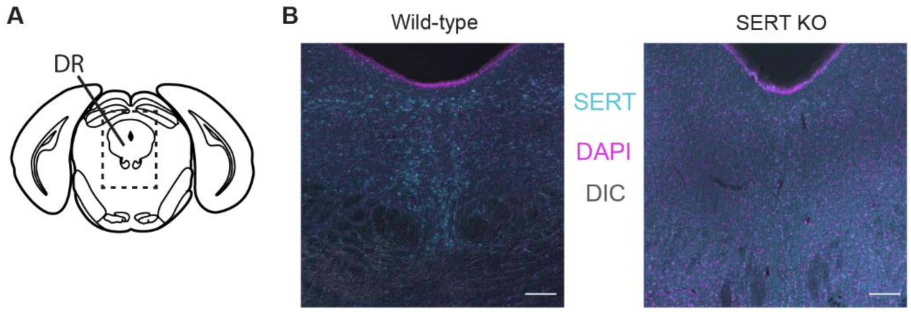

**Fig. S2. Supplementary information supporting data shown in Fig 2D. Validation of SERT antibody.** (**A**) Schematic drawing of DR, illustrating approximate antero-posterior position of slices used to test SERT antibody. (**B**) SERT antibody reveals cell bodies in the DR in a wild-type mice (*left*), but only yields background staining in a SERT KO mouse (*right*). anti-SERT, cyan; DAPI, magenta; DIC, gray. Scale bar = 100 μm.

#### Figure S3

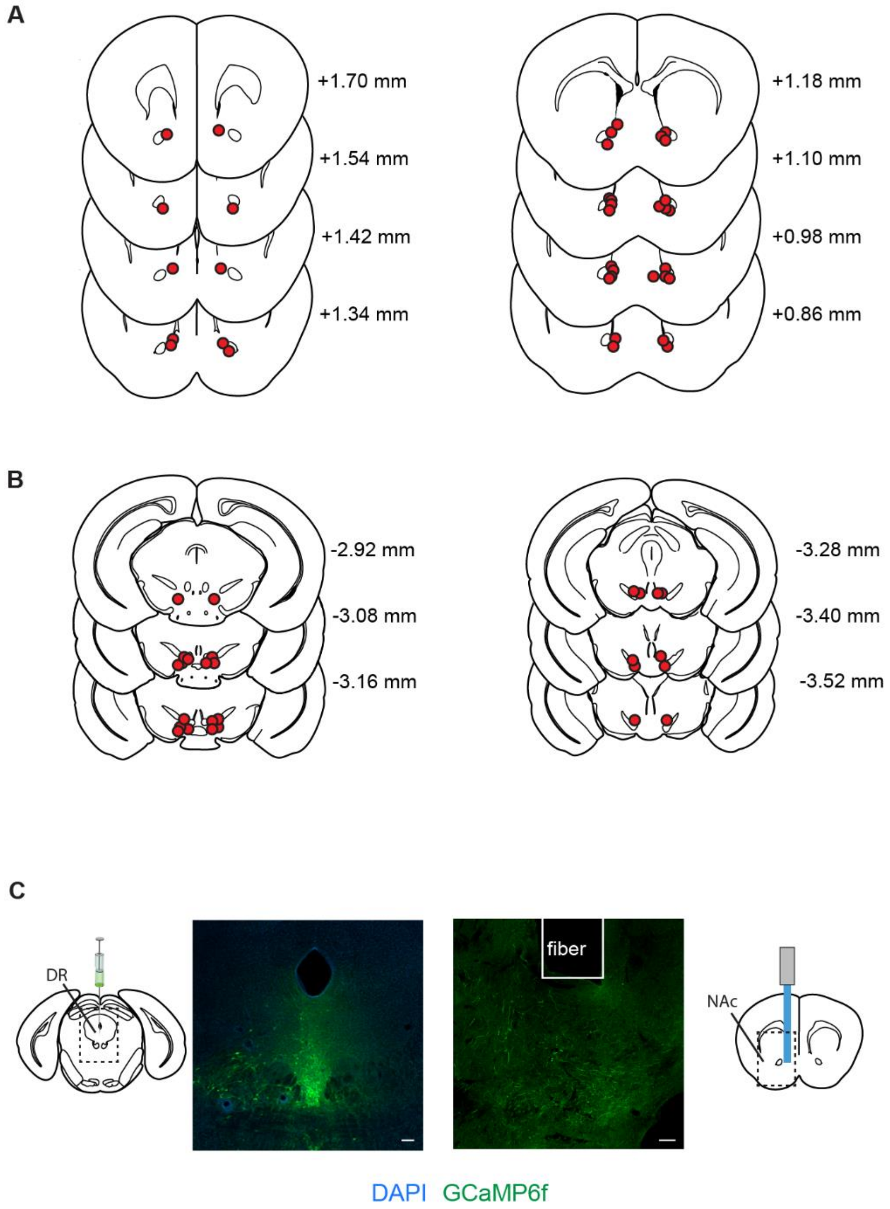

**Fig. S3. Supplementary information supporting data shown in Fig. 3.** (**A** and **B)** Location of infuser tips of cannulae implanted for intra-NAc drug delivery (**A**, Fig. 3A, C, D), and intra-VTA drug delivery (**B**, Fig. 3B). (**C**) Sample images showing virus injection (DJ-AAV-EF1a-DIO-GCaMP6f) into the DR of SERT-Cre mice *(left)* and optical fiber placement in the NAc *(right)*, for fiber photometry experiments (Fig. 3A-G). GCaMP6f-positive processes, presumed to originate from SERT+ neurons in the DR, are visible below the tip of the fiber in the NAc. GCaMP6f, green; DAPI, blue. Scale bar = 100 μm.

| Panel | Group | N | Mean ± SEM (unit) | Primary Statistic | *Post hoc* test[a] |
|---|---|---|---|---|---|
| 1B | SAL | 13 | empty, 11.92 ± 0.98 (min)<br>mouse, 12.52 ± 1.10 (min) | One-way ANOVA, unmatched;<br>$F_{7,82}$=6.20, P<0.0001 | ns |
| | MDMA 3 mg/kg | 9 | empty, 7.76 ± 1.29 (min)<br>mouse, 11.92 ± 1.67 (min) | | ns |
| | MDMA 7.5 mg/kg | 13 | empty, 7.17 ± 1.13 (min)<br>mouse, 16.21 ± 1.72 (min) | | P<0.0001 |
| | MDMA 15 mg/kg | 10 | empty, 7.94 ± 1.52 (min)<br>mouse, 15.86 ± 1.98 (min) | | P=0.0015 |
| 1C | SAL | 16 | | Two-way ANOVA, ordinary;<br>Treatment: $F_{1,163}$ = 8.42,<br>P=0.0042<br>Time: ns | |
| | MDMA 7.5 mg/kg | 14 | | | |
| 1D | SAL | 16 | -6.31 ± 7.691 (soc. ind.) | Unpaired t-test;<br>P=0.022 | |
| | MDMA 7.5 mg/kg | 14 | 24.49 ± 10.42 (soc. ind.) | | |
| 1E | SAL, both mice | 12 | -2.488 ± 11.26 (soc. ind.) | One-way ANOVA, unmatched;<br>$F_{3,61}$=5.87, P=0.0014 | --- |
| | MDMA, cup mouse | 18 | 14.75 ± 5.14 (soc. ind.) | | *vs* SAL, ns |
| | MDMA, free mouse | 20 | 26.55 ± 7.25 (soc. ind.) | | *vs* SAL, P=0.023 |
| | MDMA, both mice | 15 | 42.30 ± 5.42 (soc. ind.) | | *vs* SAL,<br>P=0.0005 |
| 1F | SAL | 10 | 2.12 ± .33 (m) | One-way ANOVA, unmatched;<br>$F_{3,38}$=9.14, P=0.0001 | --- |
| | MDMA 7.5 mg/kg | 10 | 2.18 ± .56 (m) | | *vs* SAL, ns |
| | MDMA 15 mg/kg | 11 | 5.79 ± .65 (m) | | *vs* SAL,<br>P=0.0002 |
| 1H | MDMA 7.5 mg/kg | 10 | pre, 15.96 ± 1.02<br>post, 15.75 ± 1.465 | Paired t-test;<br>ns | |
| | MDMA 15 mg/kg | 11 | pre 14.66 ± 1.45<br>post, 18.08 ± 1.43 | Paired t-test;<br>P=0.012 | |
| S1B | Female SAL | 8 | -11.1 ± 14.38 (soc. ind.) | Unpaired t-test;<br>P=0.027 | |
| | Female MDMA 7.5 mg/kg | 6 | 41.15 ± 14.36 (soc. ind.) | | |
| | Male SAL | 13 | -0.7 ± 11.26 (soc. ind.) | Unpaired t-test;<br>P=0.027 | |
| | Male MDMA 7.5 mg/kg | 13 | 40.14 ± 13.23 (soc. ind.) | | |
| S1C | SAL | 16 | -11.96 ± 10.99 (soc. ind.) | Unpaired t-test;<br>P=0.049 | |
| | MDMA | 12 | 26.99 ± 16.17 (soc. ind.) | | |
| S1D | Open arm, SAL | 8 | 2.93 ± 0.27 (min) | Unpaired t-test;<br>P=0.020 | |
| | Open arm, MDMA 7.5 mg/kg | 8 | 1.95 ± 0.26 (min) | | |
| | Closed arm, SAL | | 6.87 ± 0.27 (min) | Unpaired t-test;<br>P=0.020 | |
| | Closed arm, MDMA 7.5 mg/kg | | 7.86 ± 0.26 (min) | | |

**Table S1. Numerical and statistical data supporting Figure 1 and S1.** [a] Planned *post hoc* comparisons performed using Sidak correction for multiple comparisons.

| Panel | Group | N | Mean ± SEM (unit) | Primary Statistic | *Post hoc* test[a] |
|---|---|---|---|---|---|
| 2A | SCIT + SAL | 8 | 3.02 ± 12.78 (soc. ind.) | One-way ANOVA, unmatched; $F_{2,20}$=4.55, P=0.0235 | *vs* SAL + MDMA 7.5 mg/kg, P=0.048 |
| | SAL + MDMA 7.5 mg/kg | 9 | 42.41 ± 10.70 (soc. ind.) | | --- |
| | SCIT + MDMA 7.5 mg/kg | 6 | -4.25 ± 12.59 (soc. ind.) | | *vs* SAL + MDMA 7.5 mg/kg, P<0.030 |
| 2E | $SERT^{wt/wt}$ SAL | 22 | 1.56 ± 9.78 (soc. ind.) | Unpaired t-test; P=0.0037 | |
| | MDMA 7.5 mg/kg | 14 | 54.8 ± 14.89 (soc. ind.) | | |
| | $SERT^{wt/fl}$ SAL | 17 | 13.24 ± 11.17 (soc. ind.) | Unpaired t-test; ns | |
| | MDMA 7.5 mg/kg | 16 | 5.21 ± 17.37 (soc. ind.) | | |
| | $SERT^{fl/fl}$ SAL | 8 | 32.73 ±12.54 (soc. ind.) | Unpaired t-test; P=0.034 | |
| | MDMA 7.5 mg/kg | 14 | -28.85 ± 19.00 (soc. ind.) | | |
| 2F | $SERT^{wt/wt}$ MDMA 15 mg/kg | 10 | Pre, 14.70 ± 0.73 (min) Post, 17.14 ± 1.09 (min) | Paired t-test; P=0.038 | |
| | $SERT^{wt/fl}$ MDMA 15 mg/kg | 22 | Pre, 15.43 ± 0.36 (min) Post, 17.48 ± 0.93 (min) | Paired t-test; P=0.033 | |
| | $SERT^{fl/fl}$ MDMA 15 mg/kg | 11 | Pre, 14.75 ± 0.70 (min) Post, 16.53 ± 1.02 (min) | Paired t-test; P=0.049 | |
| 2G | SAL + SAL | 15 | -16.11 ± 13.61 (soc. ind.) | One-way ANOVA, unmatched; $F_{2,30}$=6.32, P=0.0051 | *vs* JHW + MDMA 7.5 mg/kg, P=0.0084 |
| | JHW + MDMA 7.5 mg/kg | 8 | 53.33 ± 15.06 (soc. ind.) | | --- |
| | JHW + SAL | 10 | -25.85 ± 17.33 (soc. ind.) | | *vs* JHW + MDMA 7.5 mg/kg, P=0.0055 |
| 2H | SAL | 10 | | Two-way ANOVA, ordinary; Treatment: $F_{1,163}$ = 8.42, P=0.029 Time: ns | |
| | METH 2 mg/kg | 9 | | | |
| 2I | SAL, Day 0 | 8 | 2.96 ± 0.27 (m) | One-way ANOVA, repeated measures; $F_{3,21}$=50.96, P<0.0001 | *vs* METH 2mg/kg Day 1, P=0.0010 |
| | METH 2 mg/kg, Day 1 | | 6.90 ± 0.71 (m) | | --- |
| | METH 2 mg/kg, Day 2 | | 12.29 ± 1.06 (m) | | --- |
| | METH 2 mg/kg, Day 3 | | 13.30 ± 1.06 (m) | | *vs* METH 2mg/kg Day 1, P<0.00001 |
| 2J | METH 2 mg/kg | 12 | Pre, 13.92 ± 1.57 (min) Post, 17.00 ± 1.07 (min) | Paired t-test; P=0.031 | |

**Table S2. Numerical and statistical data supporting Figure 2.** [a] Planned *post hoc* comparisons performed using Sidak correction for multiple comparisons.

| Panel | Group | N | Mean ± SEM (unit) | Primary Statistic | *Post hoc* test[a] |
|---|---|---|---|---|---|
| 3A | Intra-NAc SCIT + SAL i.p. | 20 | 11.54 ± 7.91 (soc. ind.) | One-way ANOVA, unmatched; $F_{2,51}$=3.36, P=0.043 | *vs* Intra-NAc SAL + MDMA 7.5 mg/kg i.p., ns |
| | Intra-NAc SAL + MDMA 7.5 mg/kg i.p. | 15 | 34.71 ± 13.94 (soc. ind.) | | --- |
| | Intra-NAc SCIT + MDMA 7.5 mg/kg i.p. | 19 | -6.52 ± 11.28 (soc. ind.) | | *vs* Intra-NAc SAL + MDMA 7.5 mg/kg i.p., P=0.025 |
| 3B | Intra-VTA SCIT + SAL i.p. | 18 | -7.52 ± 9.01 (soc. ind.) | One-way ANOVA, unmatched; $F_{2,41}$=7.08, P=0.0023 | *vs* Intra-VTA SAL + MDMA 7.5 mg/kg i.p., P=0.0042 |
| | Intra-VTA SAL + MDMA 7.5 mg/kg i.p. | 13 | 38.92 ± 10.38 (soc. ind.) | | --- |
| | Intra-VTA SCIT + MDMA 7.5 mg/kg i.p. | 13 | 35.67 ± 11.42 (soc. ind.) | | *vs* Intra-VTA SAL + MDMA 7.5 mg/kg i.p., ns |
| 3C | Intra-NAc SAL + MDMA 7.5 mg/kg i.p. | 13 | 40.14 ± 13.23 (soc. ind.) | One-way ANOVA, unmatched; $F_{2,33}$=7.89, P=0.0016 | *vs* Intra-NAc SAL + SAL i.p., P=0.014 |
| | Intra-NAc SAL + SAL i.p. | 11 | -1.27 ± 10.12 (soc. ind.) | | --- |
| | Intra-NAc MDMA + SAL i.p. | 12 | 55.70 ± 4.56 (soc. ind.) | | *vs* Intra-NAc SAL + SAL i.p., P=0.0010 |
| 3D | Intra-NAc SAL + MDMA 15 mg/kg i.p. | 16 | Pre, 14.82 ± 0.65 (min)<br>Post, 17.17 ± 0.72 (min) | Paired t-test; P=0.012 | |
| | Intra-NAc SCIT + MDMA 15 mg/kg i.p. | 16 | Pre, 15.00 ± 0.44 (min)<br>Post, 18.52 ± 0.92 (min) | Paired t-test; P=0.0015 | |
| | Intra-NAc RAC + MDMA 15 mg/kg i.p. | 13 | Pre, 14.86 ± 0.98 (min)<br>Post, 15.13 ± 1.16 (min) | Paired –test; ns | |
| 3G | SAL | 3 | 94.21 ± 0.40 (% baseline std. dev.) | Unpaired t-test; P<0.0001 | |
| | MDMA 7.5 mg/kg | 3 | 21.74 ± 4.46 (% baseline std. dev.) | | |
| 3K | SERT-Cre, + GCaMP6f | 5 | Mouse, 2140 ± 445.3 (cum. z)<br>Empty, 941.9 ± 263.4 (cum. z) | Unpaired t-test; P=0.0492 | |
| 3L | SERT-Cre, + GCaMP6f | 5 | Slope = 0.22 ± 0.057;<br>Y-intercept, 1.77 ± 0.41;<br>X-intercept, -5.40 | Linear regression; $F_{1,3}$=14.73, P=0.031 | |

**Table S3. Numerical and statistical data supporting Figure 3.** [a] Planned *post hoc* comparisons performed using Sidak correction for multiple comparisons.

| Panel | Group | N | Mean ± SEM (unit) | Primary Statistic | *Post hoc* test[a] |
|---|---|---|---|---|---|
| 4A | OTRA + SAL | 8 | -13.95 ± 9.84 (soc. ind.) | One-way ANOVA, unmatched; $F_{4,43}$=3.11, P=0.025 | --- |
| | SAL + MDMA 7.5 mg/kg | 12 | 32.88 ± 11.10 (soc. ind.) | | *vs* OTRA+ SAL, P=0.031 |
| | OTRA + MDMA 7.5 mg/kg | 12 | 30.70 ± 11.51 (soc. ind.) | | *vs* OTRA + SAL, P=0.042 |
| | OTRA high dose + MDMA 7.5 mg/kg | 8 | 34.24 ± 10.61 (soc. ind.) | | *vs* OTRA + SAL, P=0.046 |
| | OTRA multiple dose + MDMA 7.5 mg/kg | 8 | 46.45 ± 16.23 (soc. ind.) | | *vs* OTRA + SAL, P=0.0086 |
| 4B | Intra-NAc SAL + SAL i.p. | 11 | -10.65 ± 11.69 (soc. ind.) | One-way ANOVA, unmatched; $F_{3,35}$=7.85, P=0.0004 | --- |
| | Intra-NAc SAL + MDMA 7.5 mg/kg i.p. | 13 | 39.06 ± 11.23 (soc. ind.) | | *vs* Intra-NAc SAL + SAL i.p., P=0.024 |
| | Intra-NAc OTRA + MDMA 7.5 mg/kg i.p. | 9 | 74.31 ± 12.36 (soc. ind.) | | *vs* Intra-NAc SAL + SAL i.p., P=0.0003 |
| 4C | SAL | 17 | -2.39 ± 11.78 (soc. ind.) | Unpaired t-test; P=0.048 | |
| | MDMA 7.5 mg/kg | 14 | 33.29 ± 12.51 (soc. ind.) | | |
| 4F | D1 MSN (7 mice) | 8 | 72.78 ± 7.98 (% baseline EPSC) | Unpaired t-test, ns | |
| | D2 MSN (6 mice) | 7 | 65.84 ± 5.84 (% baseline EPSC) | | |
| 4H | MDMA (5 mice) | 8 | 70.10 ± 6.92 (% baseline EPSC) | Unpaired t-test, P=0.0087 | |
| | NAS-181 (3 mice) | 5 | 101.4 ± 5.44 (% baseline EPSC) | | |
| 4I | Intra-NAc SAL + MDMA 7.5 mg/kg i.p. | 12 | 32.85 ± 15.38 (soc. ind.) | Unpaired t-test, P=0.035 | |
| | Intra-NAc NAS-181 + MDMA 7.5 mg/kg i.p. | 11 | -17.72 ± 16.43 (soc. ind.) | | |
| 4J | SAL | 16 | -2.69 ± 10.77 (soc. ind.) | One-way ANOVA, unmatched; $F_{3,46}$=3.43, P=0.025 | --- |
| | FEN 1 mg/kg | 10 | 7.41 ± 4.01 (soc. ind.) | | *vs* SAL, ns |
| | FEN 5 mg/kg | 13 | 29.69 ± 8.27 (soc. ind.) | | *vs* SAL, P=0.048 |
| | FEN 10 mg/kg | 11 | 34.42 ± 12.04 (soc. ind.) | | *vs* SAL, P=0.027 |
| 4K | SAL | 16 | | Two-way ANOVA, ordinary; Treatment: $F_{5,138}$=19.76, P<0.0001. Time: ns | |
| | FEN 10 | 11 | | | |
| 4L | FEN (3 mice) | 5 | 69.35 ± 4.93 (% baseline EPSC) | | |

**Table S4. Numerical and statistical data supporting Figure 4.** [a] Planned *post hoc* comparisons performed using Sidak correction for multiple comparisons.
